# Mesozooplankton Distribution in the Northeastern Arabian Sea During the Spring Intermonsoon

**DOI:** 10.64898/2026.08.03.742178

**Authors:** Maria Jani Mol, Honey U. K. Pillai

## Abstract

Mesozooplankton is a vital component of pelagic food webs, serving as a key trophic link between primary producers and higher marine organisms, and driving coastal and oceanic biogeochemical cycles. This study investigated the abundance, biovolume, community structure, and spatial distribution of mesozooplankton in relation to hydrographic variables in the Northeastern Arabian Sea (NEAS 15^◦^N − 21^◦^N) during the Spring Intermonsoon (March 2019) on board *FORV Sagar Sampada*. Sampling across 14 stations comparing coastal (> 50 m) and oceanic (< 1000 m) waters revealed distinct environmental gradients.

Both Sea Surface Temperature (SST: 24.33^◦^C − 28.68^◦^C) and Sea Surface Salinity (SSS: 35.25 − 36.65) exhibited a declining trend toward northern latitudes, whereas Dissolved Oxygen (DO: 4.55 − 5.24 ml ⋅ L^−1^) increased northward. Chlorophyll *concentrations* were significantly higher in coastal regions (mean 1.45 mg ⋅ m^−3^) than in oceanic waters (mean 0.36 mg ⋅ m^−3^). Correspondingly, mesozooplankton biovolume (excluding large gelatinous taxa) was threefold higher in coastal waters (mean 1.39 mL ⋅ m^−3^) compared to oceanic waters (mean 0.38 mL ⋅ m^−3^). Mean numerical abundance was also higher coastally (605 ind. ⋅ m^−3^) than oceanically (524 ind. ⋅ m^−3^).

A total of 21 mesozooplankton taxa were identified. Copepods dominated the community structure across both regimes (coastal: 61.07%; oceanic: 56.69%). In coastal waters, Chaetognatha (14.76%), Cladocera (9.39%), and Decapod larvae (5.69%) were the secondary drivers, with localised salp and siphonophore swarms observed at 17^◦^ − 19^◦^N. In oceanic waters, Ostracoda (22.93%) and Salpidae (12.51%) were major secondary contributors. These results indicate that during the Spring Intermonsoon, the NEAS exhibits moderate productivity representative of a receding winter cooling regime.

## 1. Introduction

The global oceans cover roughly 71% of Earth’s surface and represent one of the largest carbon reservoirs on the planet, driving climate regulation, thermal absorption, and marine biodiversity. Pelagic ecosystems are fundamentally sustained by planktonic communities. While autotrophic phytoplankton convert solar energy into chemical energy, heterotrophic zooplankton regulate primary production and facilitate the transfer of energy to higher trophic levels, including commercially important fisheries, seabirds, and marine mammals.

Plankton are broadly classified by size, ranging from single-celled picoplankton to megaplankton such as scyphozoans. Among these, **mesozooplankton** (200 *μ*m − 2 mm) play a pivotal role in oceanic biogeochemical cycles, carbon export, and pelagic ecosystem stability.

The Arabian Sea is a highly dynamic ocean basin characterized by strong seasonal monsoonal forcing—the Southwest Monsoon (SWM) and Northeast Monsoon (NEM)—separated by transitional Spring and Autumn Intermonsoon periods. Winter convective cooling during the NEM enriches the upper water column of the Northern Arabian Sea with nutrients, promoting elevated biological productivity that often persists into the early Spring Intermonsoon.

Understanding the spatial distribution, biovolume, and taxonomic composition of mesozooplankton during these transitional periods is vital for assessing pelagic food web dynamics. This study evaluates the mesozooplankton community structure alongside key environmental drivers (temperature, salinity, dissolved oxygen, and chlorophyll *a*) across coastal and oceanic waters of the Northeastern Arabian Sea during the Spring Intermonsoon.

## 2. Materials and Methods

### 2.1. Study Area and Sampling Design

Sampling was conducted in March 2019 onboard the Fishery Oceanographic Research Vessel (*FORV Sagar Sampada*) across 14 designated stations in the Northeastern Arabian Sea (NEAS), spanning latitudes 15^◦^N to 21^◦^N. Stations were categorized into two distinct operational regimes:

- **Coastal waters:** Depths > 50 m
- **Oceanic waters:** Depths < 1000 m

### 2.2. Environmental Data Collection

Physical and biological oceanographic parameters were recorded at each station using a CTD profiler system (Sea-Bird Electronics):

- **Sea Surface Temperature (SST)** (^◦^C)
- **Sea Surface Salinity (SSS)**
- **Dissolved Oxygen (DO)** (ml ⋅ L^−1^)
- **Chlorophyll *a*** (mg ⋅ m^−3^)

### 2.3. Zooplankton Collection and Analysis

Mesozooplankton samples were collected using a standard **Bongo Net** equipped with a calibrated digital flowmeter to determine the volume of water filtered.

- **Biovolume Measurement:** Measured onboard via the displacement method. Large gelatinous organisms (e.g., medusae, large salps) were separated prior to biovolume determination to avoid artificial inflation of displacement values (mL ⋅ m^−3^).
- **Taxonomic Identification & Counting:** Samples were preserved in 4% buffered formalin-seawater solution. In the laboratory, samples were split using a Folsom plankton splitter, identified to the lowest possible taxonomic group under stereomicroscopes, and quantified as numerical abundance (ind. ⋅ m^−3^).

## 3. Results

### 3.1. Hydrographic Parameters

Hydrographic conditions showed clear spatial variations between coastal and oceanic transects as well as along latitudinal gradients:

- **Sea Surface Temperature (SST):** Both coastal and oceanic waters exhibited a decreasing trend toward the northern stations. Coastal SST ranged from 24.33^◦^C to 27.79^◦^C, whereas oceanic SST ranged from 25.13^◦^C to 28.68^◦^C.
- **Sea Surface Salinity (SSS):** Salinity mirrored the thermal pattern, decreasing northward. Coastal SSS varied between 35.25 and 36.62, while oceanic values ranged from 35.51 to 36.65.
- **Dissolved Oxygen (DO):** In contrast to SST and SSS, DO concentrations showed an increasing latitudinal trend northward. Coastal DO values ranged from 4.55 to 5.24 ml ⋅ L^−1^, and oceanic DO ranged from 4.79 to 5.15 ml ⋅ L^−1^.
- **Chlorophyll *a*:** Coastal waters exhibited significantly higher primary productivity, with Chlorophyll *a* averaging 1.45 mg ⋅ m^−3^ (range: 0.18 − 2.31 mg ⋅ m^−3^). Oceanic waters were comparatively oligotrophic, averaging 0.36 mg ⋅ m^−3^ (range: 0.02 − 2.02 mg ⋅ m^−3^).

**Table 1:** Summary of Hydrographic and Biological Parameters in NEAS (March 2019)

| Parameter | Coastal Waters (>50 m) Range | Coastal Mean | Oceanic Waters (<1000 m) Range | Oceanic Mean |
| --- | --- | --- | --- | --- |
| **SST ($^{\circ}\text{C}$)** | 24.33 – 27.79 | — | 25.13 – 28.68 | — |
| **SSS** | 35.25 – 36.62 | — | 35.51 – 36.65 | — |
| **DO ($\text{mL} \cdot \text{L}^{-1}$)** | 4.55 – 5.24 | — | 4.79 – 5.15 | — |
| **Chlorophyll a ($\text{mg} \cdot \text{m}^{-3}$)** | 0.18 – 2.31 | 1.45 | 0.02 – 2.02 | 0.36 |
| **Biovolume ($\text{mL} \cdot \text{m}^{-3}$)** | — | 1.39 | — | 0.38 |
| Abundance (ind. · m<sup>-3</sup>) | 82 – 1354 | 605 | 108 – 1069 | 524 |

### 3.2. Mesozooplankton Biovolume and Abundance

- **Biovolume:** Non-gelatinous mesozooplankton biovolume demonstrated a threefold enrichment in coastal waters (mean 1.39 mL ⋅ m^−3^) relative to the open ocean (mean 0.38 mL ⋅ m^−3^).
- **Numerical Abundance:** Total density was higher in coastal zones (mean 605 ind. ⋅ m^−3^; range: 82 − 1354 ind. ⋅ m^−3^) than in oceanic zones (mean 524 ind. ⋅ m^−3^; range: 108 − 1069 ind. ⋅ m^−3^).

Gelatinous blooms, specifically composed of salp swarms and siphonophores, were recorded predominantly in coastal stations between 17^◦^N and 19^◦^N.

### 3.3. Mesozooplankton Community Composition

A total of **21 mesozooplankton taxa** (comprising both holoplanktonic and meroplanktonic groups) were recorded across the study region. Nineteen (19) taxa were identified in coastal samples, while all 21 taxa were present in oceanic samples.

**Table 2:** Relative Abundance (%) of Major Mesozooplankton Groups.

| Taxa / Group | Coastal Contribution (%) | Oceanic Contribution (%) |
| --- | --- | --- |
| Copepoda | 61.07% | 56.69% |
| Ostracoda | < 1.00% | 22.93% |
| Chaetognatha | 14.76% | 2.84% |
| Salpidae | 1.37% | 12.51% |
| Cladocera | 9.39% | < 1.00% |
| Decapod Larvae | 5.69% | 1.23% |
| Lucifer spp. | 2.63% | < 1.00% |
| Stomatopod Larvae | 2.30% | < 1.00% |

- **Copepods** represented the dominant group in both coastal (61.07%) and oceanic (56.69%) assemblages.
- **Coastal secondary groups** were driven by carnivorous and filter-feeding coastal organisms: Chaetognaths (14.76%), Cladocerans (9.39%), Decapod larvae (5.69%), *Lucifer* (2.63%), and Stomatopod larvae (2.30%).
- **Oceanic secondary groups** showed a shift toward Ostracods (22.93%) and tunicates like Salps (12.51%), with reduced relative proportions of Chaetognaths (2.84%) and Decapods (1.23%).

## 4. Discussion

The physical and biological trends recorded during March 2019 capture the environmental transition characteristic of the Spring Intermonsoon in the Northeastern Arabian Sea. During this period, the intense convective mixing typical of the Northeast Monsoon begins to abate.

The lower SSTs and higher oxygen levels observed in the northern transects suggest residual signals of winter convective cooling. The elevated Chlorophyll *a* concentration in coastal stations directly supported significantly higher mesozooplankton biovolume (1.39 mL ⋅ m^−3^) and numerical abundance (605 ind. ⋅ m^−3^) compared to oceanic stations.

Copepod dominance across all stations aligns with general pelagic food web models, reaffirming their role as the primary consumer link in the region. However, the contrast in secondary group dominance highlights distinct ecological regimes:

1. **Coastal Regimes:** Characterised by high secondary production involving meroplankton (decapod and stomatopod larvae) and opportunistic groups (cladocerans, chaetognaths, and salp swarms), responding to localised nutrient availability and primary production.
2. **Oceanic Regimes:** Characterised by a transition toward deeper-dwelling or pelagic specialists, evidenced by the marked increase in Ostracoda (22.93%) and pelagic tunicates.

Overall, the hydrographic conditions and plankton metrics indicate that during March 2019, the Northeastern Arabian Sea was in a receding phase of winter cooling, supporting moderate overall biological productivity.

## 5. Conclusion

This study provides key baseline insights into the mesozooplankton community structure and environmental interactions in the Northeastern Arabian Sea during the Spring Intermonsoon. The findings confirm strong coastal-oceanic and latitudinal gradients in physical parameters, primary productivity, and secondary biomass. Coastal regions served as productivity hotspots driven by elevated Chlorophyll *a* level, higher mesozooplankton biovolume, and rich meroplanktonic diversity, while oceanic regimes were characterised by high copepod and ostracod dominance.

## Acknowledgments

The authors express gratitude to the Director, Zoological Survey of India (ZSI), Kolkata, and the Ministry of Earth Sciences (MoES) / CMLRE for providing shipboard facilities aboard *FORV Sagar Sampada*. Thanks are also extended to the Faculty of Ocean Science and Technology, Kerala University of Fisheries and Ocean Studies (KUFOS), for institutional support.

---

FORV Sagar Sampada

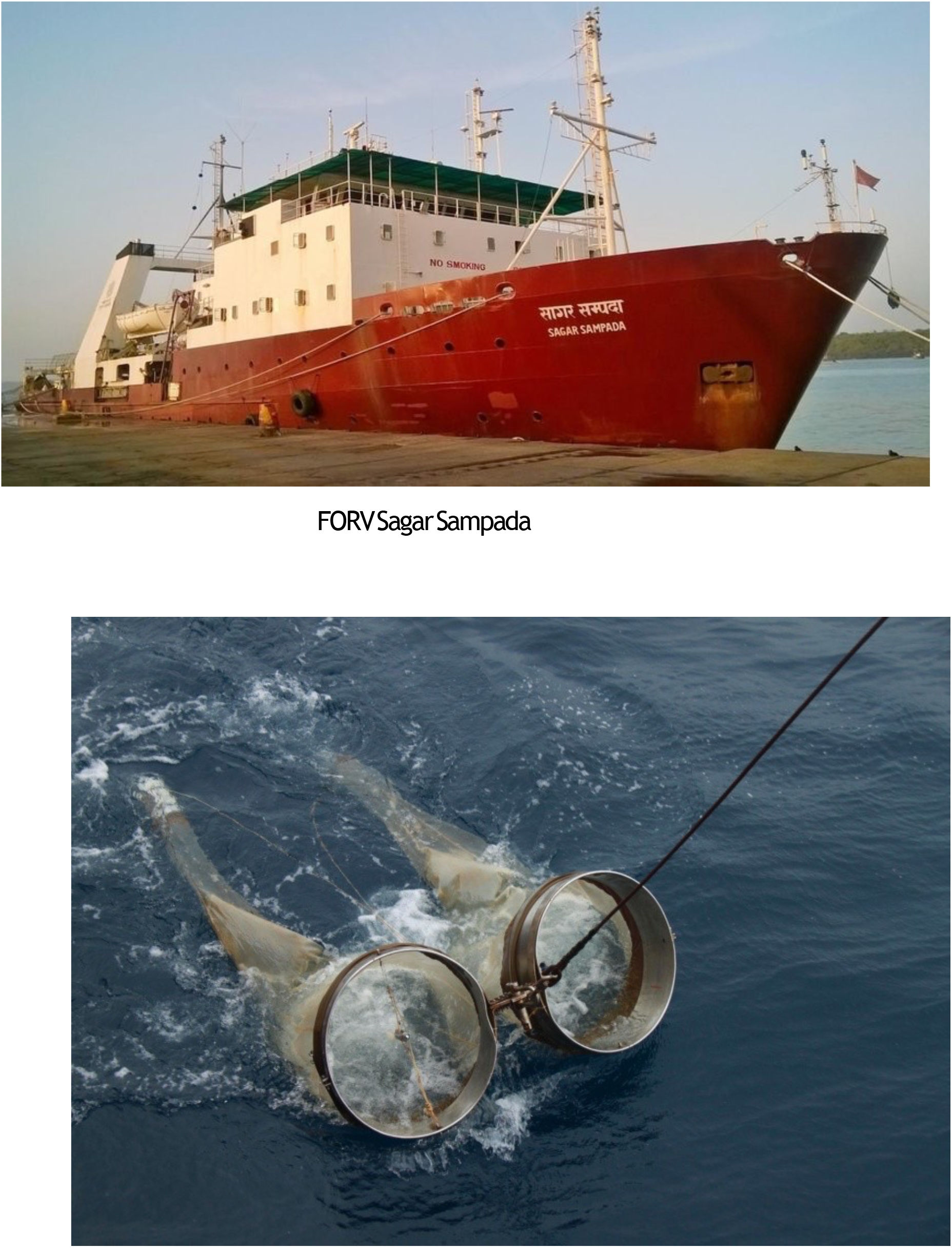

Plate 2. Bongo Net operation

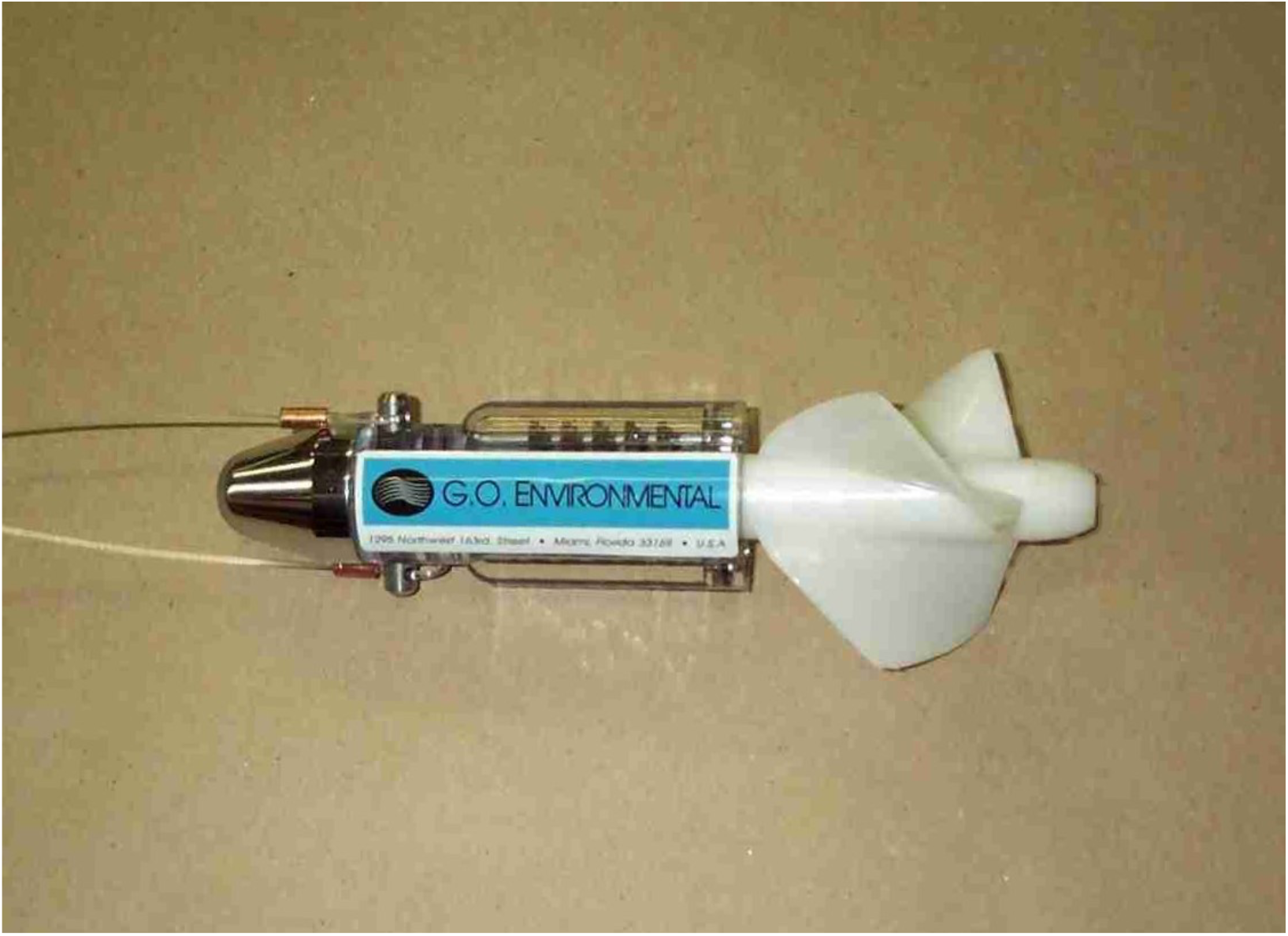

Plate 3. Flow meter

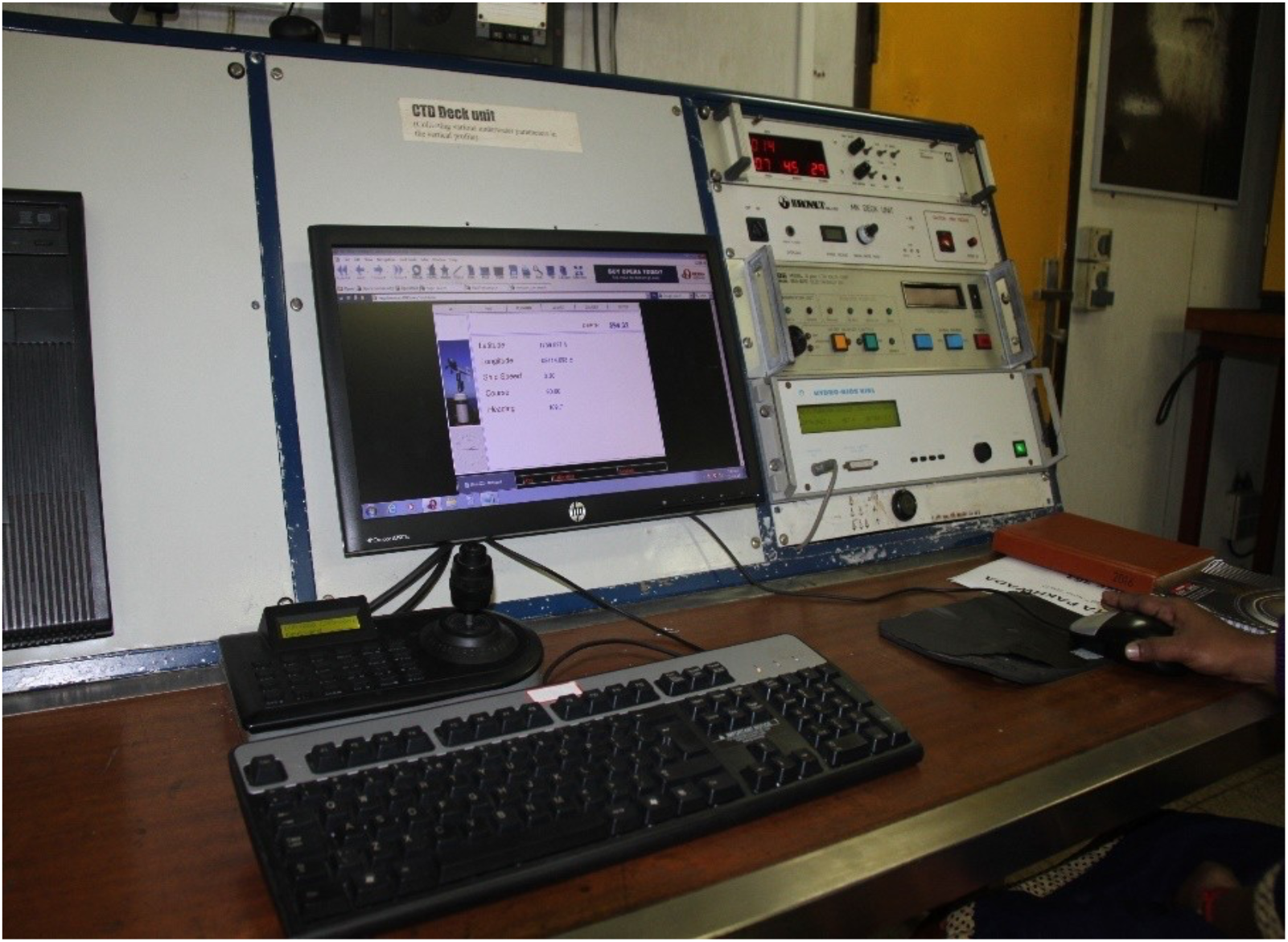

Plate 4.CTD Deck Control unit

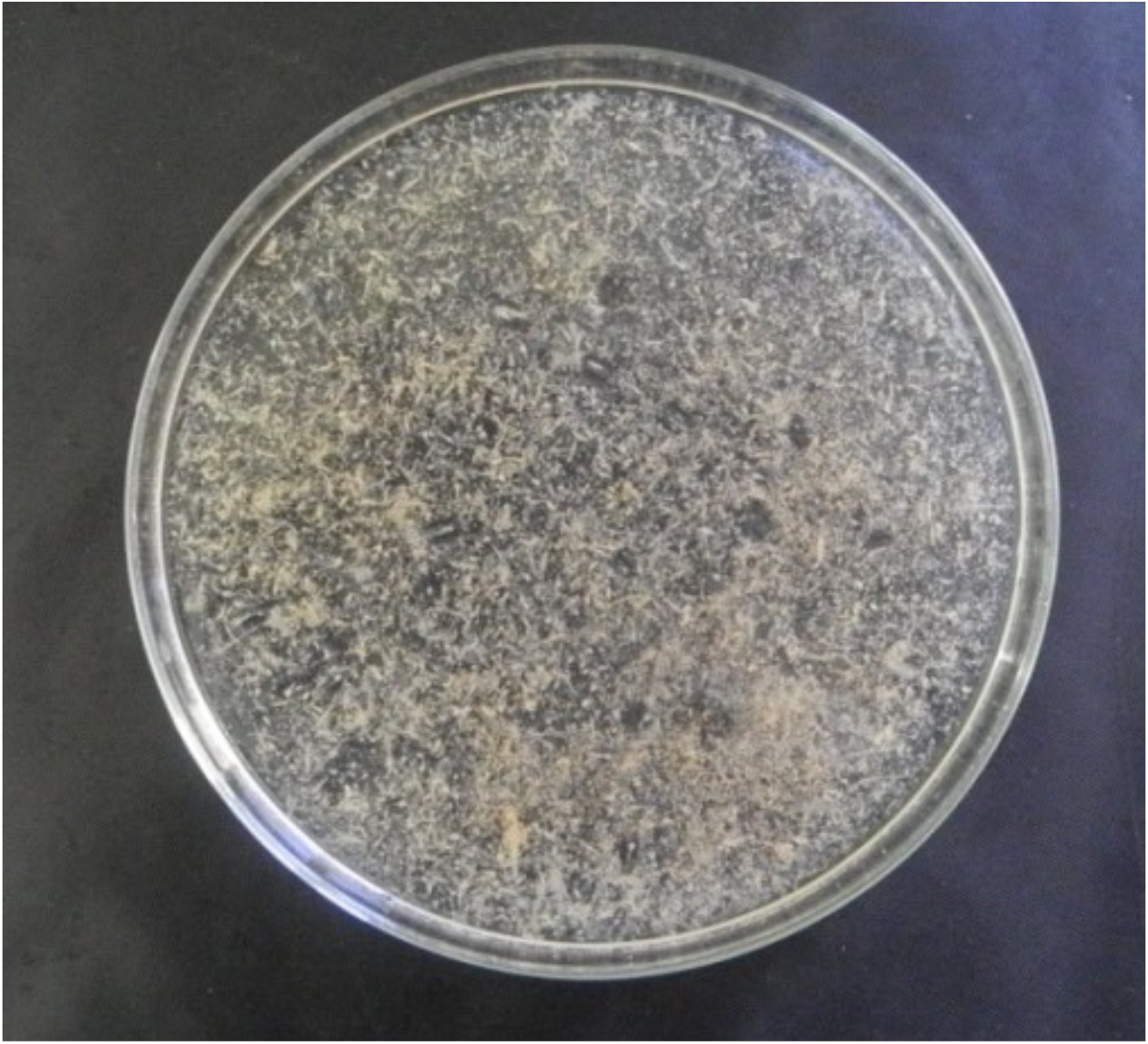

mesozooplankton assemblage

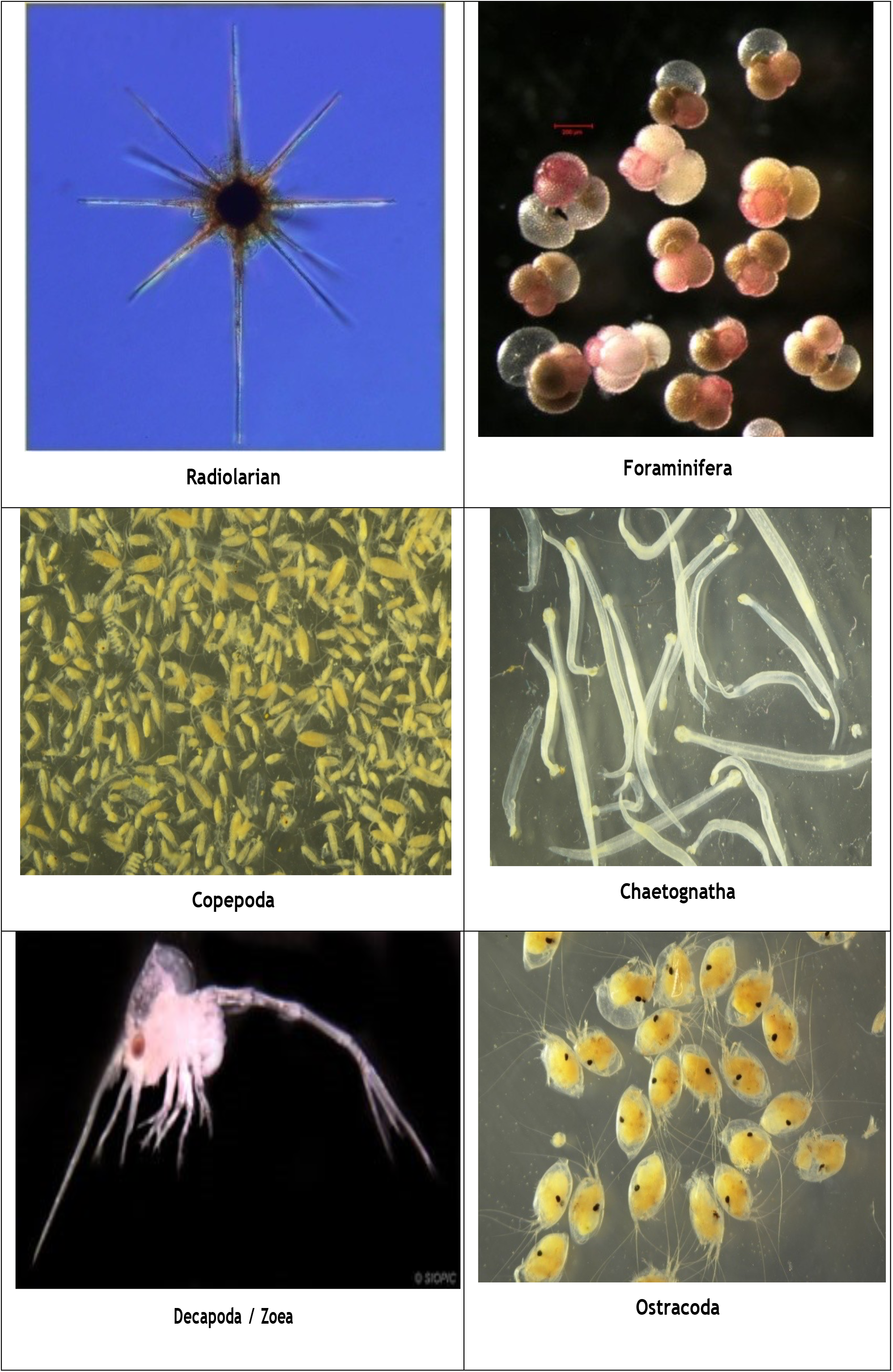
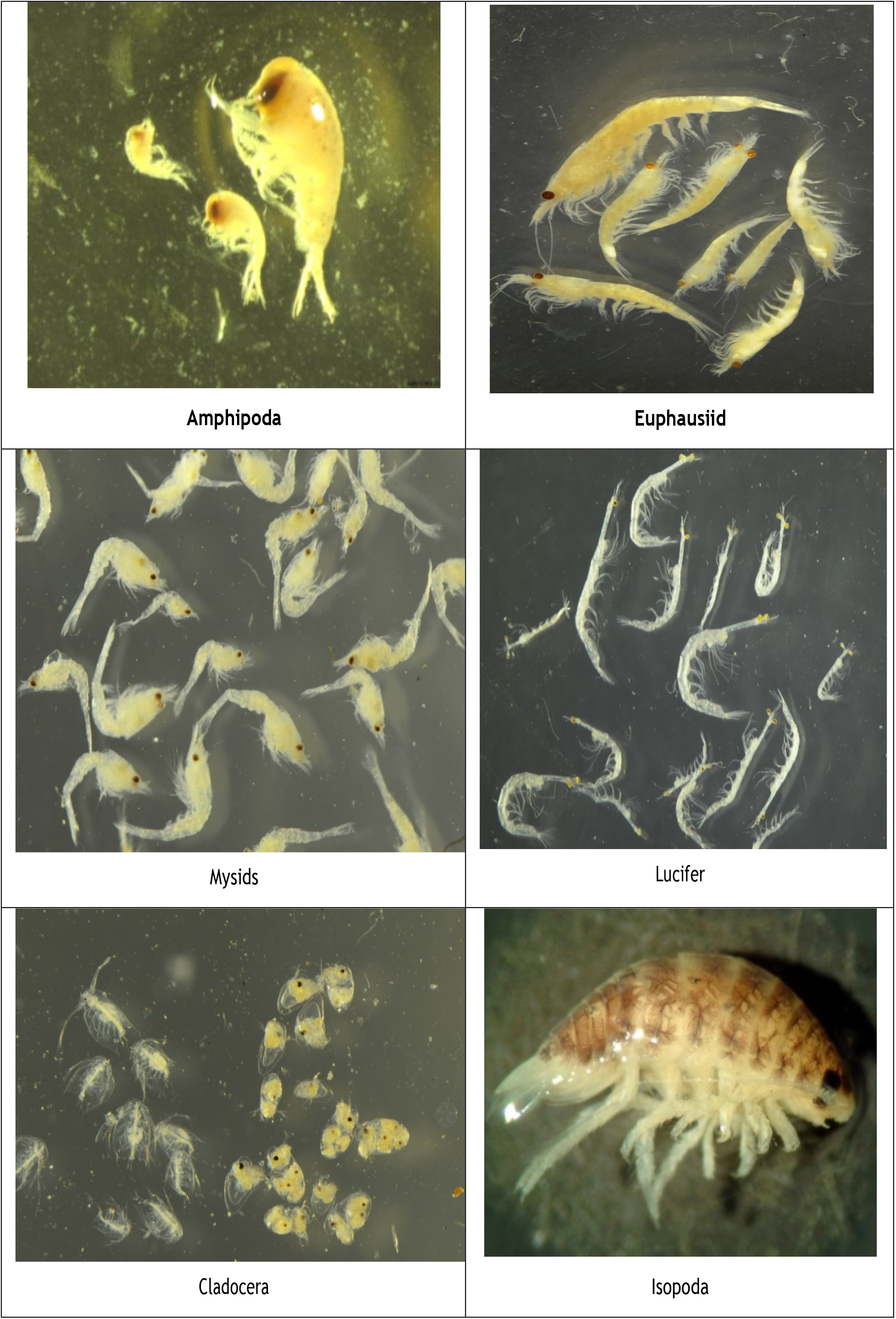
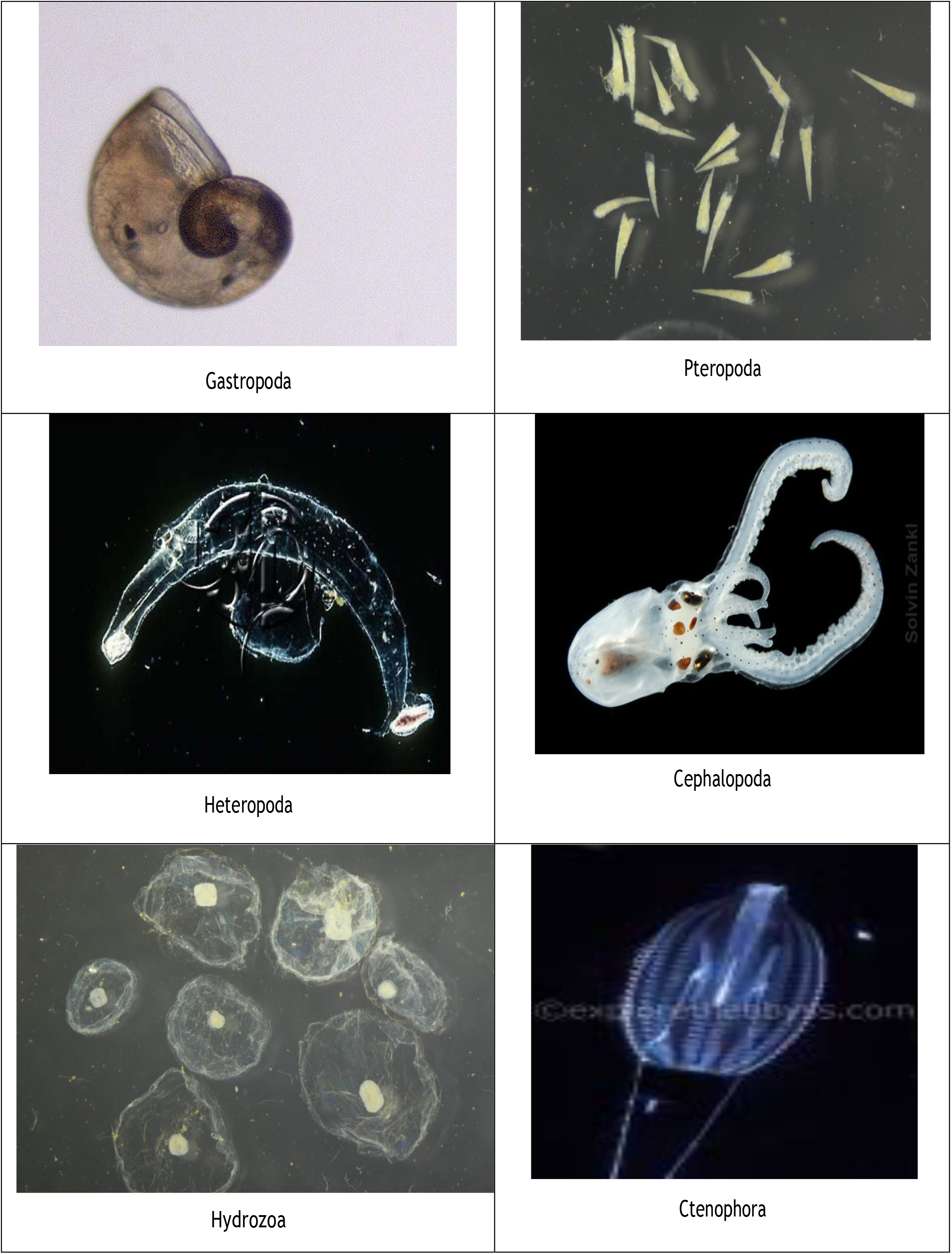
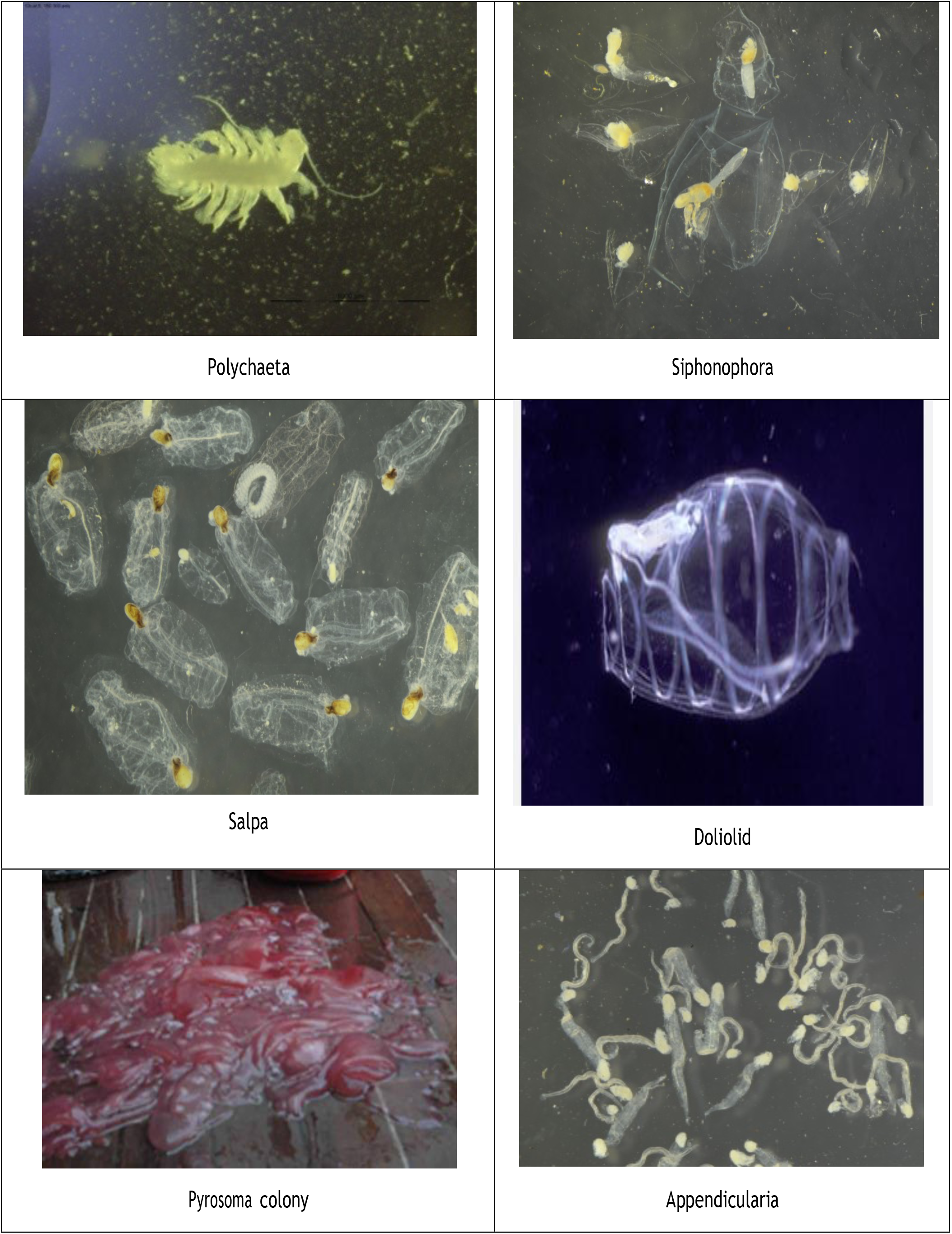
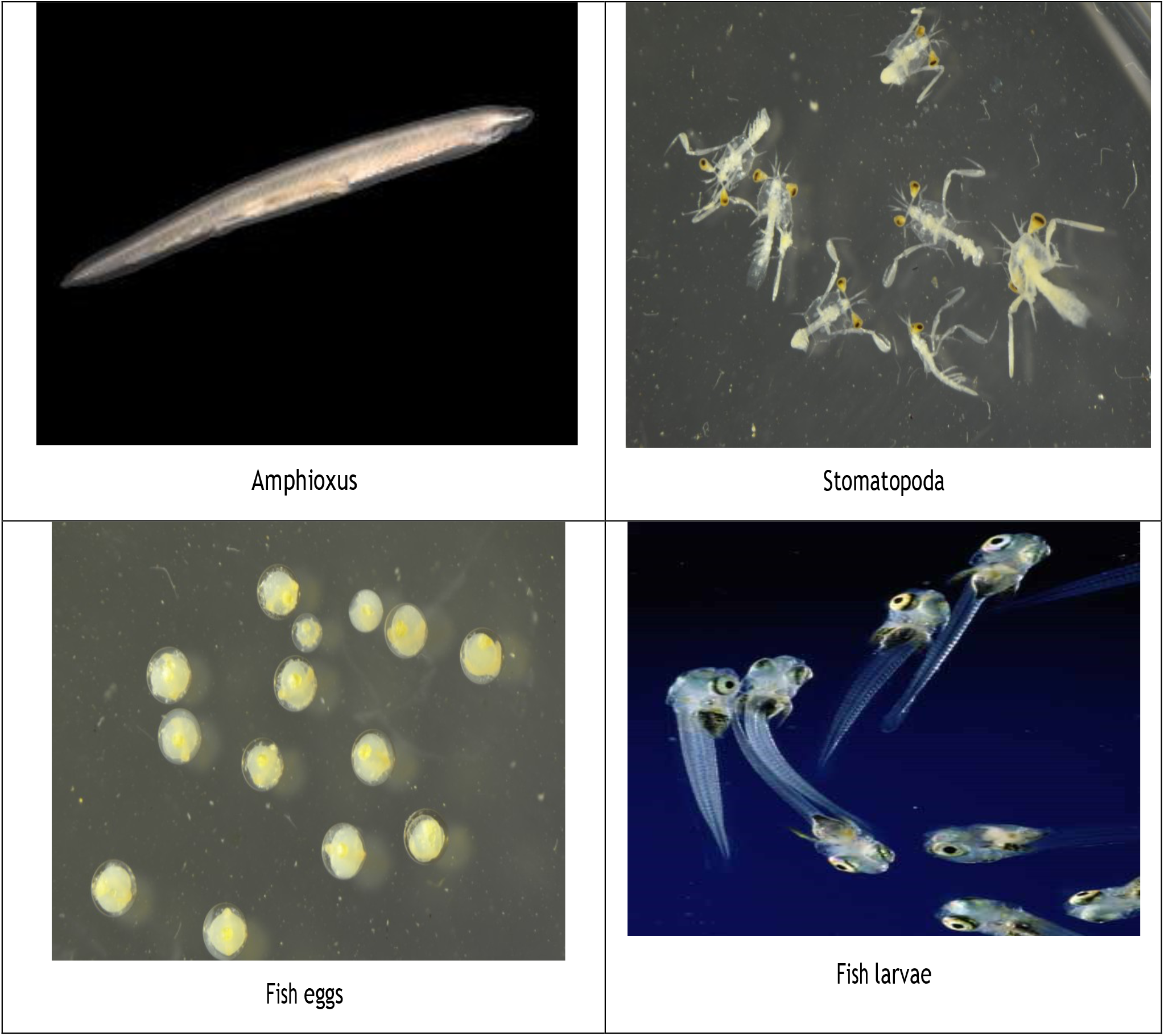

Different mesozooplankton groups

